# Genotype-free demultiplexing of pooled single-cell RNA-seq

**DOI:** 10.1101/570614

**Authors:** Jun Xu, Caitlin Falconer, Quan Nguyen, Joanna Crawford, Brett D. McKinnon, Sally Mortlock, Alice Pébay, Alex W. Hewitt, Anne Senabouth, Stacey Andersen, Nathan Palpant, Han Sheng Chiu, Grant W. Montgomery, Joseph Powell, Lachlan Coin

## Abstract

A variety of experimental and computational methods have been developed to demultiplex samples from pooled individuals in a single-cell RNA sequencing (scRNA-Seq) experiment which either require adding information (such as hashtag barcodes) or measuring information (such as genotypes) prior to pooling. We introduce scSplit which utilises genetic differences inferred from scRNA-Seq data alone to demultiplex pooled samples. scSplit also extracts a minimal set of high confidence presence/absence genotypes in each cluster which can be used to map clusters to original samples. Using a range of simulated, merged individual-sample as well as pooled multi-individual scRNA-Seq datasets, we show that scSplit is highly accurate and concordant with demuxlet predictions. Furthermore, scSplit predictions are highly consistent with the known truth in cell-hashing dataset. We also show that multiplexed-scRNA-Seq can be used to reduce batch effects caused by technical biases. scSplit is ideally suited to samples for which external genome-wide genotype data cannot be obtained (for example non-model organisms), or for which it is impossible to obtain unmixed samples directly, such as mixtures of genetically distinct tumour cells, or mixed infections. scSplit is available at: https://github.com/jon-xu/scSplit

## Background

Using single-cell RNA sequencing (scRNA-Seq) to cell biology at cellular level provides greater resolution than ‘bulk’ level analyses, thus allowing more refined understanding of cellular heterogeneity. For example, it can be used to cluster cells 7 into sub-populations based on their differential gene expression, so that different fates of cells during development can be discovered. Droplet-based scRNA-Seq (for 9 example Drop-Seq [1] or 10X Genomics Systems [2]) allows profiling large numbers of cells for sequencing by dispersing liquid droplets in a continuous oil phase [3] in an automated microfluidics system, and as a result is currently the most popular approach to scRNA-Seq despite a high cost per run. Methods that lower the per sample cost of running scRNA-Seq are required in order to scale this approach up to population scale. An effective method for lowering scRNA-Seq cost is to pool samples prior to droplet-based barcoding with subsequent demultiplexing of sequence reads.

Cell hashing [4] based on Cite-seq [5] is one such experimental approach to demultiplex pooled samples. This approach uses oligo-tagged antibodies to label cells prior to mixing, but use of these antibodies increases both the cost and sample preparation time per run. Moreover it requires access to universal antibodies for organism of interest, thus limiting applicability at this stage to human and mouse. Alternatively, computational tools like demuxlet [6] have been developed to demultiplex cells from multiple individuals, although this requires additional genotyping information to assign individual cells back to their samples of origin. This limits the utility of demuxlet, as genotype data might not be available for different species; biological material may not be avaliable to extract DNA; or the genetic differences between samples might be somatic in origin.

Another issue for droplet-based scRNA-Seq protocols is the presence of doublets, which occurs when two cells are encapsulated in same droplet and acquire the same barcode. The proportion of doublets increases with increasing number of cells barcoded in a run. It is imperative that these are flagged and removed prior to downstream analysis. Demuxlet [6] uses external genotype information to address this issue, and other tools have been developed to solve this issue based on expression data alone, including Scrublet [7] and Doubletdetection [8].

Here we introduce a simple, accurate and efficient tool, mainly for droplet-based scRNA-Seq, called “scSplit”, which uses a hidden state model approach to demultiplex individual samples from mixed scRNA-Seq data with high accuracy. Our approach does not require genotype information from the individual samples to demultiplex them, which also makes it suitable for applications where genotypes are unavailable or difficult to obtain. scSplit uses existing bioinformatics tools to identify putative variant sites from scRNA-seq data, then models the allelic counts to assign cells to clusters using an expectation-maximisation framework.

## Methods

(All relevant source codes available at https://github.com/jon-xu/scSplit/)

### Overview

The overall pipeline for the scSplit tool includes seven major steps (Figure 1):

1. Data quality control and filtering: The mixed sample BAM file is first filtered to keep only the reads with a list of valid barcodes to reduce technical noise. Additional filtering is then performed to remove reads that meet any of the following: mapping quality score less than 10, unmapped, failing quality checks, secondary or supplementary alignment, or PCR or optical duplicate. The BAM file is then marked for duplication, sorted and indexed.
2. SNV calling (Figure 1A): Freebayes v1.2 [9] is used to call SNVs on the filtered BAM file, set to ignore insertions and deletions (indels), multi-nucleotide polymorphisms (MNPs) and complex events. A minimum base quality score of one and minimum allele count of two is required to call a variant. The output VCF file is further filtered to keep only SNVs with quality scores greater than 30.
3. Building allele count matrices (Figure 1B): The “build matrices.py” script is run which produces two .csv files, one for each of reference and alternate allele counts as output.
4. Model initialization (Figure 1C): find the distinct groups of cells in the scRNA-Seq and use them to initialize the Allele Fraction Model (SNVs by samples).
5. E-M iterations till convergence (Figure 1D): Initialized allele fraction model and the two allele count matrices are used together to calculate the probability of each cell belonging to the clusters. After each round, allele fraction model is updated based on the probability of cell assignment and this is iterated until overall likelihood of the model reaches convergence.
6. Alternative presence/absence genotypes (Figure 1E): matrix indicating cluster genotypes at each SNV is built in this step.
7. Find distinguishing variants for clusters and use to assign samples to clusters (Figure 1F): In order to assign each model cluster back to the specific sample, distinguishing variants are identified so that genotyping of the least number of loci using the a suitable platform may be performed. Gram-Schmidt orthogonalization [10] is used to get the minimum set of informative P/A genotypes.

**Figure 1.**
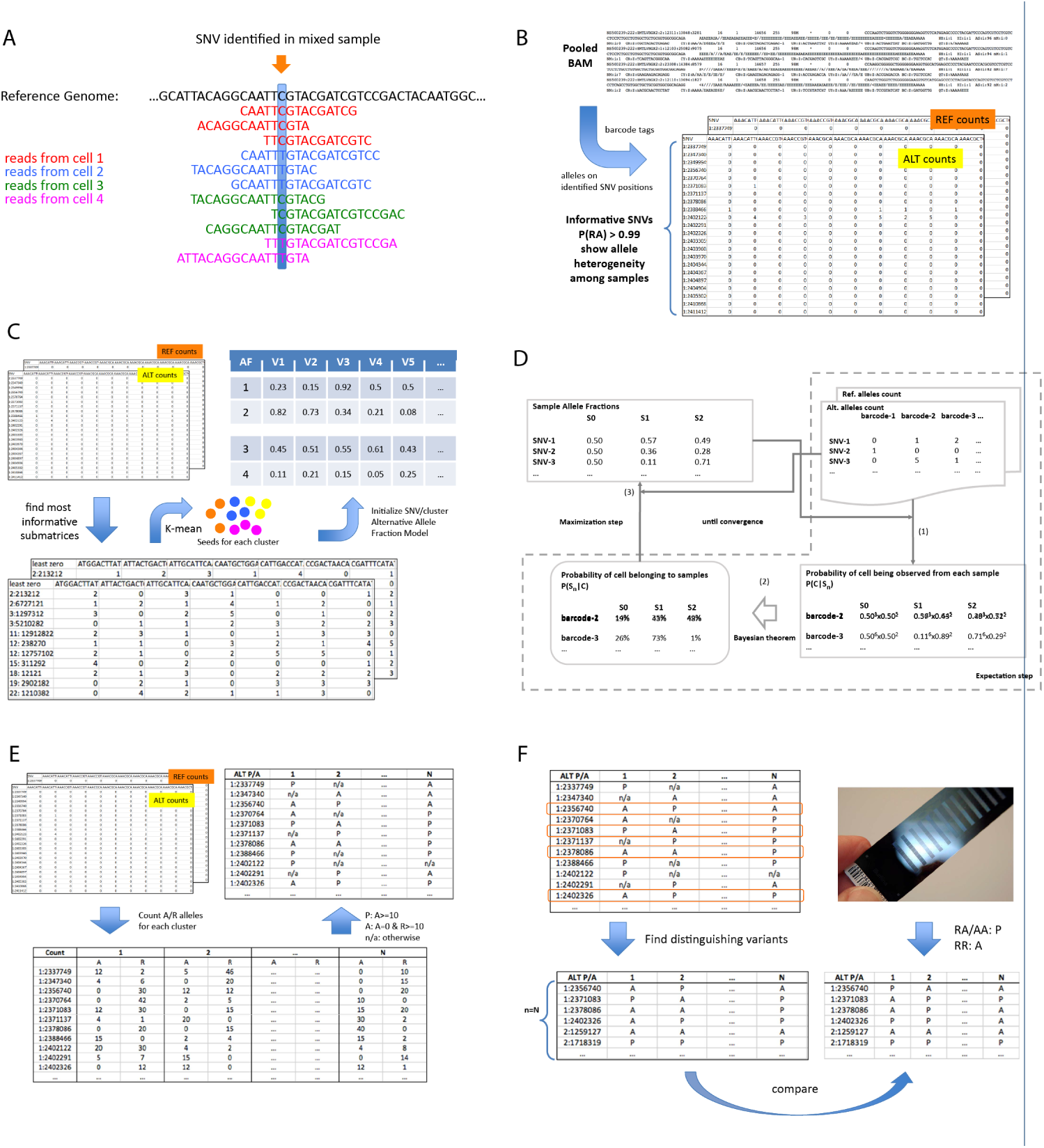
The overall pipeline of scSplit tool. (A) SNV identified based on reads from all cells which have similar or different genotypes; (B) Alternative and reference allele count matrices built from each read in the pooled-sequenced BAM at the identified informative SNVs; (C) Initial allele fraction model constructed from the initial cell seeds and their allele counts; (D) Expectation-maximization process to find the most optimized allele fraction model, based on which the cells are assigned to clusters; (E) Presence/Absence matrix of alternative alleles generated from the cell assignments; (F) minimum set of distinguishing variants found to be used to map clusters with samples.

### Data quality control

Samtools was used to filter the reads with verified barcodes for mapping and alignment status, mapping quality, and duplication (samtools view -S -bh -q 10 -F 3844 [input] >[output]). Duplicates were removed (samtools rmdup [input] [output]) followed by sorting and indexing.

### SNV calling on scRNA-Seq dataset

SNVs were called on the scRNA-Seq mixed sample BAM file with freebayes [9], a widely used variant calling tool. The freebayes arguments “-iXu -q 1” were set to ignore indels and MNPs and exclude alleles with supporting base quality scores of less than one. This generated a VCF file containing all SNVs from the mixed sample BAM file. Common SNPs of a population (for example The international Genome sample resource (IGSR)) could be used to filter the found SNVs, and minimize potential noises from non-common SNPs.

### Building allele count matrices

Allele count matrices were then built from 1) the provided mixed sample BAM file and 2) the VCF file obtained from the SNV calling program. Two allele count matrices were generated, one for the reference alleles and one for the alternate alleles, each with SNVs in rows and barcodes in columns. Each data element in the matrix indicated either the number of reference or alternate alleles detected in one cell barcode at that specific SNV position. This provided a full map of the distribution of reference and alternate alleles across all barcodes at each SNV.

The allele count matrices captured information from all reads overlapping SNVs to reflect the different allele fraction patterns from different barcodes or samples. To build the allele count matrices, pysam fetch [11] was used to extract reads from the BAM file. The reads overlapping each SNV position were fetched and counted for the presence of the reference or alternate allele. In order to increase overall accuracy and efficiency, SNVs whose GL(RA) (likelihood of heterozygous genotypes) was lower than log_10_(1 − *error*) where error = 0.01 were filtered out. These were more homozygous and thus less informative for detecting the differences between the multiple samples. The generated matrices were exported to csv files for further processing.

### Model initialization by using maximally informative cluster representatives

To initialize the model, initial probabilities of observing an alternative allele on each SNV position in each cluster were calculated. The overall matrix was sparse and a dense sub-matrix with a small number of zero count cells was generated. To do that, cells were first sorted according to their number of zero allele counts (sum of reference and alternative alleles) at all SNVs and SNVs were similarly sorted according to their number of zero allele counts (sum of reference and alternative alleles) across all cells. Next, we selected and filtered out 10% of the cells among those with the most number of zero expressed SNVs and 10% of the SNVs among those where the most number cells had zero counts. This was repeated until all remaining cells had more than 90% of their SNVs with non-zero allele counts and all SNVs had non-zero counts in more than 90% of cells. This subset of matrices was the basis for the seed barcodes to initialize the whole model. The sub-matrix was transformed using PCA with reduced dimensions and then K-means clustering was performed to split the cell subset into expected number of clusters. By using the allele fractions on the subset of SNVs in these initially assigned cells, each cluster of the model could be initialized. Let *N* (*A*_*c,v*_) and *N* (*R*_*c,v*_) be the Alternative and Reference allele counts on SNV v and cell c accordingly, and let *pseudo*_*A*_*R* be the pseudo allele count for both Alternative and Reference alleles, and *pseudo*_*A*_ be the pseudo allele count for Alternative alleles, we calculated *P* (*A*_*v*_|*S*_*n*_), the probability of observing Alternative allele on SNV v in Sample n, according to below equation:

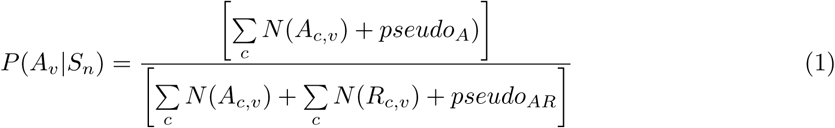

### Expectation–maximization approach

The Expectation-maximization (EM) algorithm [12] was used to conduct iterations using the full allele count matrices (Figure 1). Each iteration consisted of an E-step to calculated the probablity of seeing cells in all clusters, based on the allele fraction model, and an M-step to use the new probability of seeing cells in all clusters to update the allele fraction model. EM iterations stopped when convergence was reached, so that the overall probability of observing the cells, or the reference/alternative alleles count matrices, was maximized.

During the E-step, the tool first calculated *P* (*C*_*i*_|*S*_*n*_), the likelihood of observing a cell *C*_*i*_ in sample *S*_*n*_, which was equal to the product of the probability of observing the allele fraction pattern over each SNV, which in turn equaled to the product of probability of having observed the count of alternative alleles and probability of having observed the count of reference alleles. Let *c*_*i*_ be the i-th cell, *S*_*n*_ be the n-th sample, *A*_*v*_ be the Alternative allele on SNV v, and N(A), N(R) be the quantity of Alternative and Reference alleles:

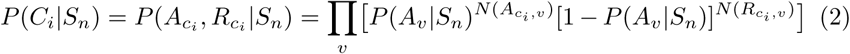

And then *P* (*C*_*i*_|*S*_*n*_) was transformed to *P* (*S*_*n*_|*C*_*i*_), the cell-sample probability, i.e. the probability of a cell *C*_*i*_ belonging to sample *S*_*n*_, using Bayes’ theorem, assuming equal sample prior probabilities (*P* (*S*_1_) = *P* (*S*_2_) = … = *P* (*S*_*n*_)):

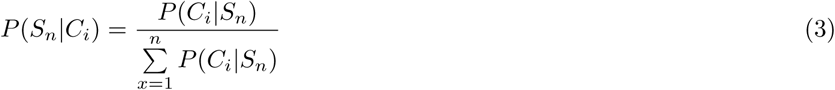

Next, weighted allele counts were distributed to the different cluster models according to the cell-sample probability, followed by the M-step, where the allele fraction model represented by the alternative allele fractions was updated using the newly distributed allele counts, so that allele fractions at all SNV positions in each sample model was recalculated:

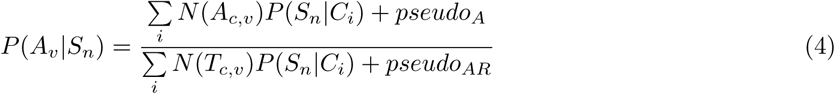

Let *P* (*S*_*n*_) be the probability of seeing the n-th sample, the overall log-likelihood of the whole model [13] was calculated as:

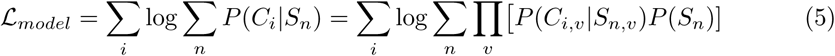

### Multiple runs to avoid local maximum likelihood

The entire process was repeated for 30 rounds with the addition of randomness during model initialization and the round with the largest sum of log likelihood was taken as the final result. Randomness was introduced by randomly selecting the 10% of cells and SNVs to be removed from the matrices during initialization from a range of the lowest ranked cells and SNVs as detailed previously.

### Cell cluster assignment

Next, probability of a cell belonging to a cluster *P* (*S*_*n*_|*C*_*i*_) was calculated. Cells were assigned to a cluster based on a minimum threshold of P>0.99. Those cells with no *P* (*S*_*n*_|*C*_*i*_) larger than the threshold were regarded as unassigned.

### Handling of doublets

During scRNA-Seq experiments, a small proportion of droplets can contain cells from more than one sample. These so called doublets, contain cells from same or different samples sharing the same barcode, which if not addressed would cause bias. Our model took these doublets into consideration. During our hidden state based demultiplexing approach, we included an additional cluster so that doublets could be captured. To identify which cluster in the model was the doublet cluster in each round, the sum of log-likelihood of cross assignments was checked:

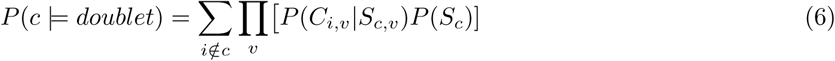

The sum log-likelihood of cells from all other clusters being assigned to a specific cluster was calculated for each cluster in turn and compared. The cluster with the largest sum log-likelihood of cross assignment was designated as the doublet cluster. We allow user input on the expected proportion of doublets. If the expected number of doublets was larger than those detected in the doublet cluster, cells with largest read depth were moved from singlet clusters to doublet cluster, so that the total number of doublets meet expectation as input.

### Alternative allele Presence/Absence genotyping for clusters

To identify a minimum set of variants, which can distinguish between sample clusters, we generated alternative allele P/A genotype matrix (SNVs by clusters). To do that, sum of reference and alternate allele counts across all cells assigned to each cluster were calculated. And for each SNV and each cluster, “P” was marked if there were more than 10 alternative allele counts, and “A” for more than 10 reference allele counts but no alternative allele count. “NA” was set if neither criterion was met.

### Mapping clusters back to individual samples using minimal set of P/A genotypes

Based on the P/A matrix, we started from informative SNVs which had variations of “P” or “A” across clusters and avoid picking those with “NA”s. Then, unique patterns involved in those SNVs were derived and for each unique P/A pattern, one allele was selected to subset the whole matrix. Next, Gram-Schmidt orthogonalization [10] was applied on the subset of P/A matrix, in order to find the variants which can be basis vectors to effectively distinguish the clusters. If not enough SNVs were found to distinguish all the clusters, the clusters were split into smaller groups so that for each group there was enough variants to distinguish the clusters within that group. And to distinguish clusters from different smaller groups, if the selected variants could not be used to distinguish any pair of clusters, additional variants were selected from the whole list of variants where no NAs were involved and P/A was different between the pair of clusters. Ideal situation was N variants for N clusters, but it was possible that >N variants were needed to distinguish N clusters.

As such, the P/A genotyping of each cluster, on the minimum set of distinguishing variants, could be used as a reference to map samples to clusters. After running genotyping on this minimum set of loci for each of the individual samples, a similar matrix based on sample genotypes could be generated, by setting the alternative presence flag when genotype probability (GP) was larger than 0.9 for RA or AA, or absence flag when GP was larger than 0.9 for RR. By comparing both P/A matrices, we could link the identified clusters in scSplit results to the actual individual samples.

In practice, samples can be genotyped only on the few distinguishing variants, so that scSplit-predicted clusters can be mapped with individual samples, while the whole genotyping is not needed. When the whole genotyping is available, we also provide an option for users to generate distinguishing variants only from variants with R2 > 0.9, so that they can compare the distinguishing matrix from scSplit with that from known genotypes on more confident variants.

### Data simulation

To test the consistency of the model, and the performance of our demultiplexing tool, reference/alternative count matrices were simulated from a randomly selected scRNA-Seq BAM file and a multi-sample VCF file. We assume the randomly selected BAM file had a representative gene expression profile.

First, data quality was checked and the BAM and VCF files were filtered. Second, barcodes contained in the BAM file were randomly assigned to samples in the VCF file, which gave us the gold-standard of cell-sample assignments to check against after demultiplexing. Then, all the reads in the BAM file were processed, that if a read overlapped with any SNV position contained in the merged VCF file, its barcode was checked to get its assigned sample and the probability *P*(*A*_*c,v*_) of having the alternative allele for that sample was calculated using the logarithm-transformed genotype likelihood (GL) or genotype probability (GP) contained in the VCF file. The probability of an allele being present at that position could then be derived so that the ALT/REF count at the SNV/barcode in the matrices could be simulated based on the alternative allele probability. Let ℒ(*AA*) and ℒ(*RA*) be the likelihood of seeing AA and RA of a certain cell c on a certain SNV v:

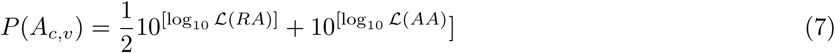

Finally, doublets were simulated by merging randomly picked 3% barcodes with another 3% without overlapping in the matrix. This was repeated for every single read in the BAM file.

With the simulated allele fraction matrices, the barcodes were demultiplexed using scSplit and the results were compared with the original random barcode sample assignments to validate. Code for simulation is available in the Github repository (https://github.com/jon-xu/scSplit).

### Single cell RNA-seq data used in testing scSplit

In table 3 and table 4, we used published hashtagged data from GSE108313 and PBMC data from GSE96583. For table 2 and table 5, endometrial stromal cells cultured from 3 women and fibroblast cells cultured from 38 healthy donors over the age of 18 years respectively were run through the 10x Genomics Chromium 3’ scRNA-seq protocol. The libraries were sequenced on the Illumina Nextseq 500. FASTQ files were generated and aligned to Homo sapiens GRCh38p10 using Cell Ranger. Individuals were genotyped prior to pooling using the Infinium PsychArray.

**Table 1.** Overview of accuracy and performance of scSplit on simulated mixed samples, with one CPU and 30GB RAM. We used PBMC donor B [2] and genotype data from demuxlet [6] as simulation templates

| Simulation | sim2 | sim3 | sim4 | sim8 |
| --- | --- | --- | --- | --- |
| Mixed samples | 2 | 3 | 4 | 8 |
| Number of cells | 12,383 | 12,383 | 12,383 | 12,383 |
| Reads per cell | 1,041 | 1,041 | 1,041 | 1,041 |
| Informative SNVs | 34,116 | 34,116 | 34,116 | 34,116 |
| Assign cells | 41min | 41min | 46min | 47min |
| Singlet TPR | 0.97 | 0.97 | 0.97 | 0.97 |
| Singlet FDR | 0 | 9E-5 | 9E-5 | 9E-5 |
| Doublet TPR | 0.997 | 0.997 | 0.997 | 0.997 |
| Doublet FDR | 0 | 0 | 0 | 0 |

**Table 2.** Comparison of scSplit and demuxlet performance in demultiplexing merged three individually genotyped stromal samples (TPR: True Positive Rate; FDR: False Discovery Rate); Total cell numbers: 9,567; Reads per cell: 14,495; Informative SNVs: 63,129; Runtime for matrices building: 67 min, Runtime for cell assignment: 55 min

| Predictions vs Truth |  | TPR | FDR |
| --- | --- | --- | --- |
| scSplit | Singlet | 0.94 | 0.02 |
|  | Doublet | 0.65 | 0.04 |
| demuxlet | Singlet | 0.93 | 0.02 |
|  | Doublet | 0.66 | 0.47 |

**Table 3.** Comparison of scSplit and demuxlet performance in demultiplexing hashtagged and multiplexed eight individually genotyped PBMC samples (TPR: True Positive Rate; FDR: False Discovery Rate); Total cell numbers: 7,932; Reads per cell: 5,835; Informative SNVs: 16,058; Runtime for matrices building: 35 min, Runtime for cell assignment: 20 min

| Predictions vs Truth |  | TPR | FDR |
| --- | --- | --- | --- |
| scSplit | Singlet | 0.96 | 0.13 |
|  | Doublet | 0.35 | 0.28 |
| demuxlet | Singlet | 0.79 | 0.10 |
|  | Doublet | 0.65 | 0.46 |

**Table 4.** Comparison of scSplit and demuxlet performance in demultiplexing multiplexed eight individually genotyped PBMC samples (TPR: True Positive Rate; FDR: False Discovery Rate); Total cell numbers: 6,145; Reads per cell: 33,119; Informative SNVs: 22,757; Runtime for matrices building: 45 min, Runtime for cell assignment: 35 min

| scSplit vs demuxlet | TPR | FDR |
| --- | --- | --- |
| Singlets | 0.80 | 0.18 |
| Doublets | 0.12 | 0.92 |

**Table 5.**
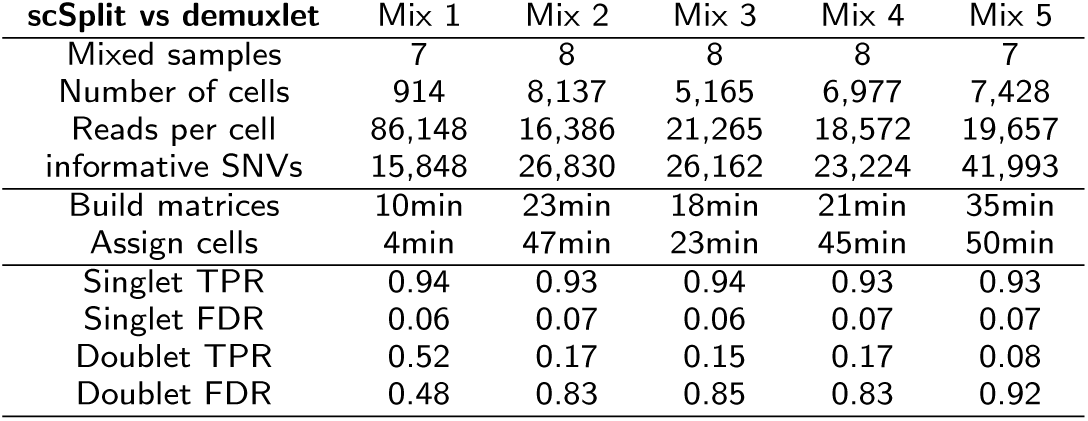
Overview of accuracy and performance running scSplit on five multiplexed scRNA-Seq datasets, with one CPU and 30GB RAM. (TPR: True Positive Rate; FDR: False Discovery Rate)

## Results

Our new tool, scSplit for demultiplexing pooled samples from scRNA-Seq data, only requires the FASTQ files obtained from single cell sequencing, together with a white-list of barcodes, while it does not require either genotype data, nor a list of common variants. All result data are available via https://github.com/jon-xu/scSplit_paper_data.

### Runs simulated on two-, three-, four- and eight-mixed samples shows high accuracy and efficiency of scSplit

We used a single scRNA-seq BAM file from Zheng et al. [2] as a template for simulation. Additionally, we took eight unrelated samples (1043, 1249, 1511, 1493, 1598, 1085, 1079, 1154) from a merged VCF file used in figure 2 supplementary data of Kang et al. [6], as the source of multi-sample genotype likelihoods for simulation (see “Data simulation” in Methods). We ran simulation tests using our scSplit tool and used the distinguishing variants to identify the individual donor for each cluster. In order to assess the accuracy of the method, we calculated both the proportion of cells from each cluster which were correctly assigned to it among the true correct number of cells in each cluster (True Positive Rate or TPR), as well as the proportion of cells assigned to a cluster which were incorrect against the total assigned cells (False Discovery Rate or FDR). We also report the average TPR and average FDR. We obtained very high overall TPR (0.97) and low FDR (less than 1e-4) for two-, three-, four- and eight-mixed samples, and also very accurate doublet predictions (Table 1). All clusters including doublet cluster match perfectly with known samples (Figure 2A).

**Figure 2.**
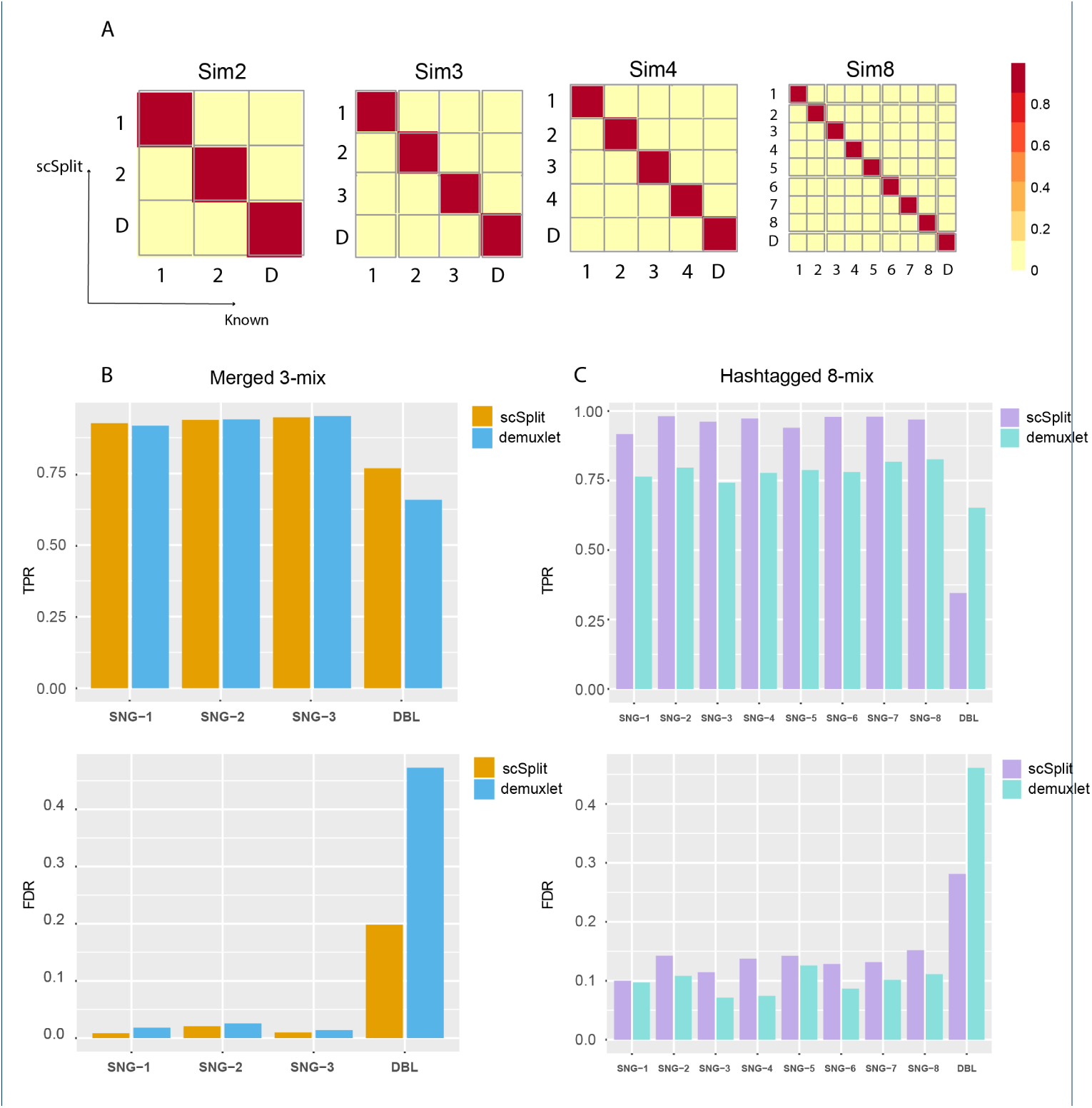
Results on simulated, merged, pooled RNA-Seq datasets confirmed scSplit a useful tool to demultiplex pooled single cells. (A) Confusion matrix showing scSplit demultiplexing results on simulated 2-, 3-, 4- and 8-mix; (B) TPR and FDR of for singlets and doublets predicted by scSplit and demuxlet compared to known truth before merging; (C) TPR and FDR of for singlets and doublets predicted by scSplit and demuxlet compared to cell hashing tags.

### scSplit performs similarly well as demuxlet in demultiplexing merged individually-sequenced three stromal samples

We then tried running scSplit on a manual merging of three individually-sequenced samples. We merged the BAM files from three individual samples (Methods). In order to create synthetic doublets, we randomly chose 500 barcodes whose reads were merged with another 500 barcodes. We ended up with 9,067 singlets and 500 doublets, knowing the their sample origins prior to merging. Both scSplit and demuxlet [6] pipelines were run on the merged samples, and the results were compared with the known truth. We observed high concordance of singlets prediction between both tools (TPR/FDR: 0.94/0.02 vs 0.93/0.02), and a better doublet prediction from scSplit compared to demuxlet (TPR/FDR: 0.65/0.04 vs 0.66/0.47) (Figure 2B and Table 2).

### scSplit predictions highly consistent with known source of hashtagged and pooled eight PBMC samples

Next, we tested scSplit on a published scRNA-Seq dataset (GSE108313) which used cell-hashing technology to mark samples of the cells before multiplexing [4]. We ran through the scSplit pipeline with the SNVs filtered by common SNVs provided on The International Genome Sample Resource (IGSR) [**?**].

According to the scSplit pipeline, distinguishing variants were identified, and P/A matrix was generated to assign the cells to clusters (Methods). We then extracted the reference and alternative allele absence information at these distinguishing variants from the sample genotypes and generated a similar P/A matrix. Both matrices were compared so that clusters were mapped to samples (Figure S1).

Our results were highly consistent with the known cell hashing tags (Table 3). We saw higher TPR for singlets in scSplit (0.96) than demuxlet (0.79) and similar singlet FDRs (0.10 vs 0.13). Although the doublet TPR of scSplit (0.35) was lower than that of demuxlet (0.65), the doublet FDR (0.28) was better than demuxlet (0.46). If the expected number of doublets was selected higher, cells with largest read depth could be moved from singlet clusters to doublet cluster to increase the TPR for doublets with a decrease of TPR for singlets.

We also compared the performance of overall P/A genotyping matrices generated based on scSplit and demuxlet predictions against that from the known genotypes (“Alternative allele Presence/Absence genotyping for clusters” section of Methods). The results show that genotypes inferred from both scSplit and demuxlet predictions have good concordance with sample genotypes (Table S1).

### Comparing scSplit with demuxlet on more pooled scRNA-Seq samples

We ran scSplit on published data from demuxlet paper [6] (Figure 3A and Table 4). By taking demuxlet predictions as ground truth, we achieved high singlet TPR (0.80), although the doublet prediction of the two tools were quite distinct to each other.

**Figure 3.**
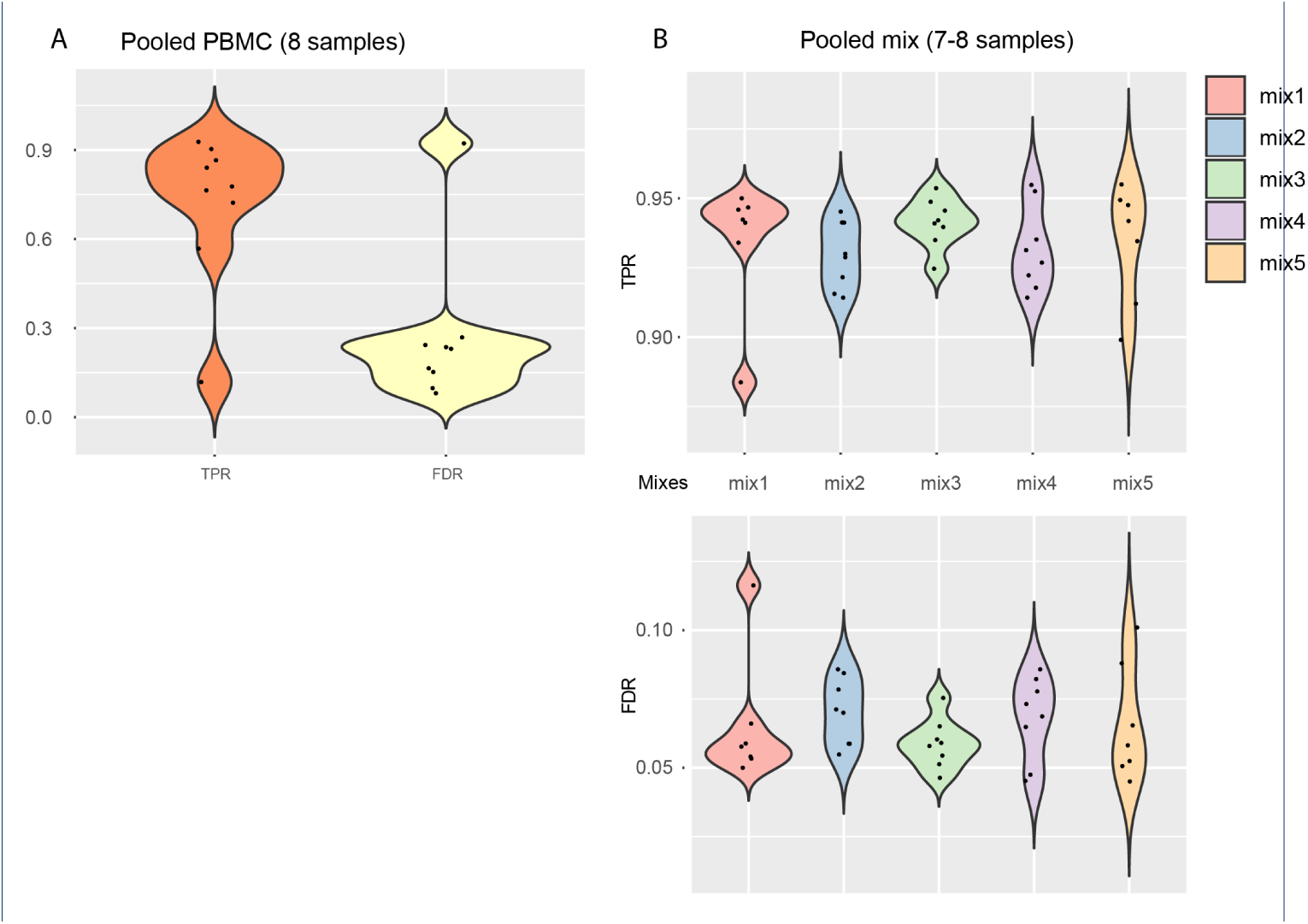
Results of scSplit on pooled PBMC scRNA-Seq and that on a set of pooled fibroblast samples. (A) TPR and FDR compared to demuxlet predictions on pooled PBMC scRNA-Seq; (B) Violin plot of singlet TPR and FDR for five 7- or 8-mixed samples based on scSplit vs demuxlet.

We also ran our tool on a set of genotyped and then pooled fibroblast scRNA-Seq datasets. Predictions from scSplit and demuxlet showed high concordance in singlets prediction (TPR: 0.93-0.94, FDR: 0.06-0.07), although not on doublets (TPR: 0.08-0.52, FDR: 0.45-0.92) when demuxlet was treated as gold-standard (Figure 3B and Table 5). Mapping between clusters and samples were recorded in Figure S2.

### Pooled samples show less batch effects than individually sequenced ones

We further checked the gene expression profiles of the three individual samples mentioned previously (Figure 2B and Table 2), in two different scenarios: 1) sequenced individually; 2) pooled the samples followed by demultiplexing after sequencing.

The results were plotted using Uniform Manifold Approximation and Projection for Dimension Reduction (UMAP [14]). The three samples were clearly separated when individually sequenced and not normalized (Figure 4A), while the samples were normalized using scran [15], we observed a different distribution of samples (Figure 4B). We saw similar pattern when we plotted the samples which were pooled and sequenced together, even without normalization (Figure 4C). And the normalization had less effect in the pooled scenario in a pooled situation (Figure 4D). In general, we showed a clear batch effect when samples were sequenced in different runs, which could be minimized if they were multiplexed before sequencing. Pooling multiple samples and sequence them together followed by demultiplexing processes could minimize the need for normalization and keep as much information as possible for downstream analyses.

**Figure 4.**
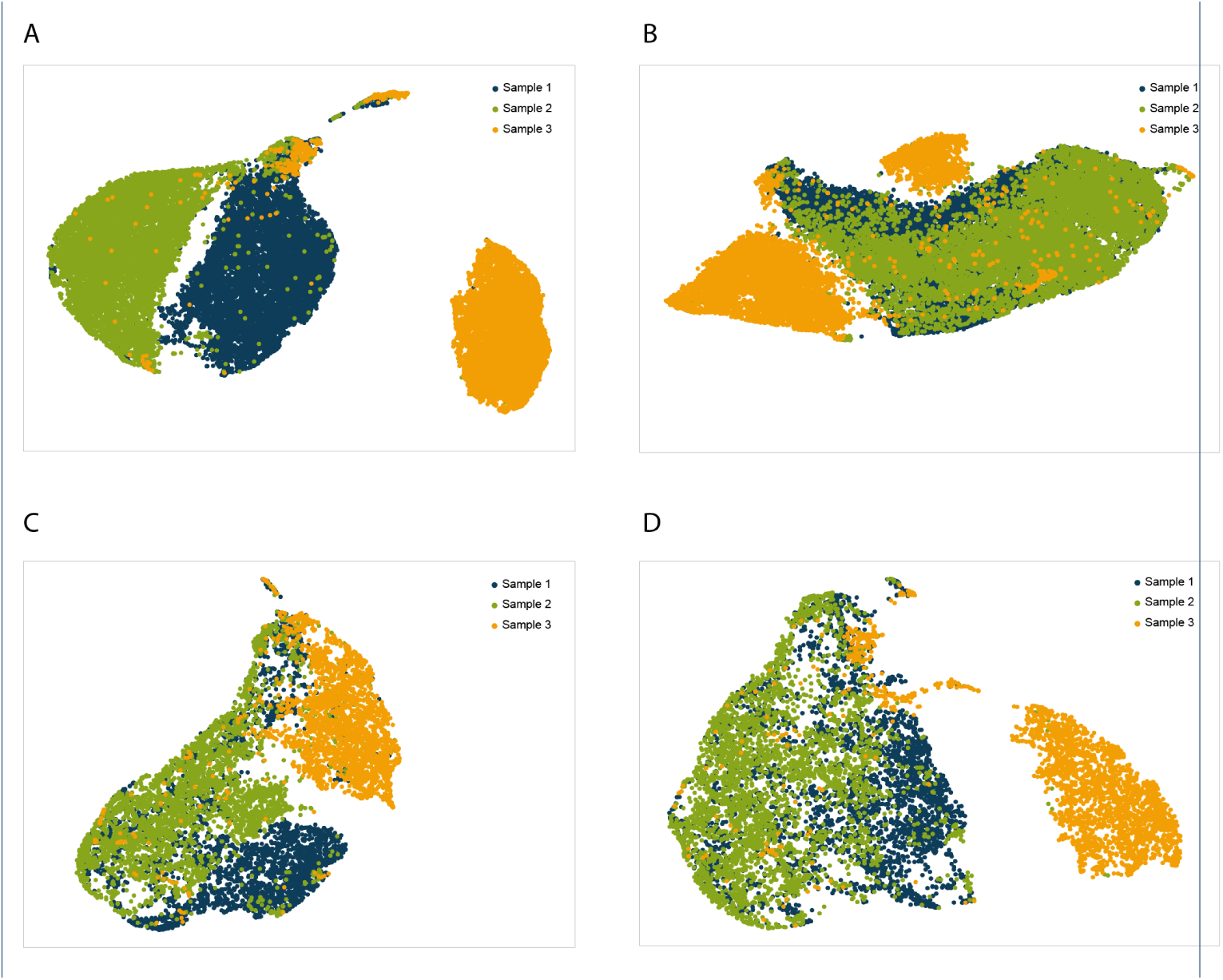
Batch effect during sequencing runs found in comparison of individually was obvious compared to pooled RNA-Seq data. (A) UMAP for three individually sequenced samples; (B) UMAP for three individually sequenced and normalized samples; (C) UMAP for pooled sequencing of same three individual samples, samples marked based on demultiplexing results using scSplit; (D) UMAP for pooled sequencing of same three individual samples, normalized by total sample reads

## Discussion

We developed the scSplit toolset to facilitate accurate, cheap and fast demultiplexing of mixed scRNA samples, without needing sample genotypes prior to mixing. scSplit also generates a minimum set of alleles (as few as the sample numbers), enabling researchers to link the resulting clusters with the actual samples by comparing the allele presence at these distinguishing loci. When predefined individual genotypes are not available as a reference, this can be achieved by designing a simple assay focused on these distinguishing variants (such as a Massarray or multiplexed PCR assay). Although the tool was mainly designed for droplet-based scRNA-Seq, it can also be used for scRNA-Seq data generated from other types of scRNA-Seq protocols.

We filtered out indels, MNPs and complex rearrangements when building the model and were able to show that SNVs alone provide adequate information to delineate the differences between multiple samples. As an alternative to using allele fractions to model multiple samples, genotype likelihoods could also be used for the same purpose, however more memory and running time would likely be needed, especially when barcode numbers in mixed sample experiments increase. Our tests showed no discernable difference in accuracy between these two methods.

The current version of scSplit assumes that the number of mixed samples are known. It is possible to run the scSplit tool for different sample numbers and compare the model log-likelihoods to select the most likely number of samples being modelled, but this would require significant computational resources and time. Alternatively, the reference and alternative allele counts in different samples and the size of the doublet cluster could be used to determine the sample number. Further optimization of the tool would be needed to effectively implement these options. Although scSplit was mainly tested on human samples, it can also be applied to other organisms, and is especially useful for those species without dense genotyping chips available. We also expect the application of scSplit in cancer related studies, to distinguish tumor cells from healthy cells, as well as to distinguish tumor sub clones.

## Conclusions

scSplit is an accurate, efficient and computationally resource-wise method with which to conduct demultiplexing of individual cells from pooled samples of scRNA-Seq. Our method and software is broadly applicable in a variety of biological and medical research areas, including but not limited to, distinguishing mixed infections, delineating tumor sub-clones and sequence analysis in non-model organisms.

## Acknowledgement

We thank Rahul Satija and Shiwei Zheng for providing helpful data from CITE-Seq based hashtagged scRNA-Seq study [4]. We also thank fundings from: National Health and Medical Research Council Career Development Fellowship (LC, APP1130084), National Health and Medical Research Council (NHMRC), Practitioner Fellowship (AWH), Senior Research Fellowship (AP), Australian Research Council Future Fellowship (AP, FT140100047), Stem Cells Australia – the Australian Research Council Special Research Initiative in Stem Cell Science (JEP, AWH, AP, NP)

## Author contributions statement

LC and CF initiated the project. LC and JX designed the algorithms. JX implemented the toolset in Python and tested it on multiple datasets. JP and BM own sample datasets. AP and AH generated the fibroblast bank. NP and HC cultured the cells in fibroblast samples. QN, SM and AS preprocessed the sample datasets. JP, QN, GM, BM, SM, JC and SA participated in important discussions and provided useful suggestions on different issues. JX drafted the manuscript, LC, JP, GM, QN and JC reviewed and revised the manuscript.

## Declarations

I can confirm I have included a statement regarding data and material availability in the method section of my manuscript.

---

**Table S1.** Accuracy of alternative allele Presence/Absence genotypes built from scSplit/demuxlet clusters compared with that from sample genotyping, based on Hashtag scRNA-Seq dataset. Accuracy is the proportion of correctly generated genotypes among all non-NA genotypes; NA is the percentage of NA genotypes among all possible elements (shared SNVs x samples)

| Genotyping | Accuracy | NA |
| --- | --- | --- |
| scSplit | 0.712 | 77% |
| demuxlet | 0.725 | 79% |

**Figure S1.**
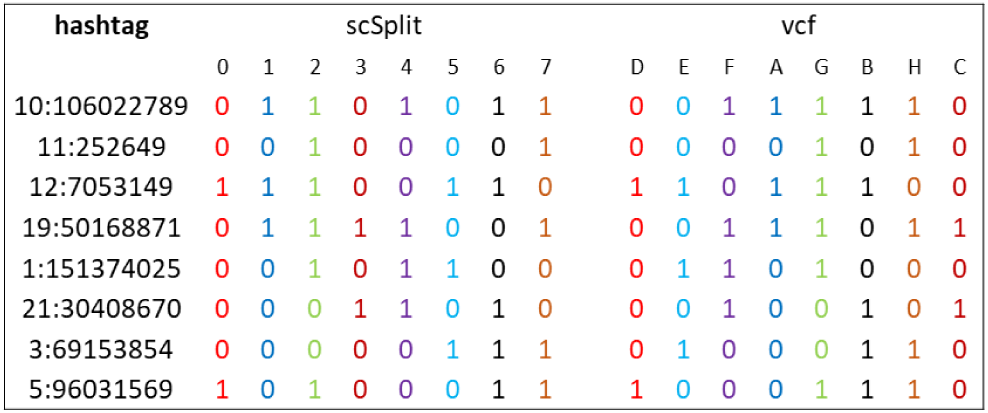
Illustration of presence absence matrices calculated on pooled scRNA-seq datasets. Within each rectangle we show the P/A genotype for each sample caclulated on scRNA-Seq data using scSplit, and the corresponding P/A genotype calculated from matched genotype files. The mapping of scRNA-Seq sample to genotype sample is indicated by colors.

**Figure S2.**
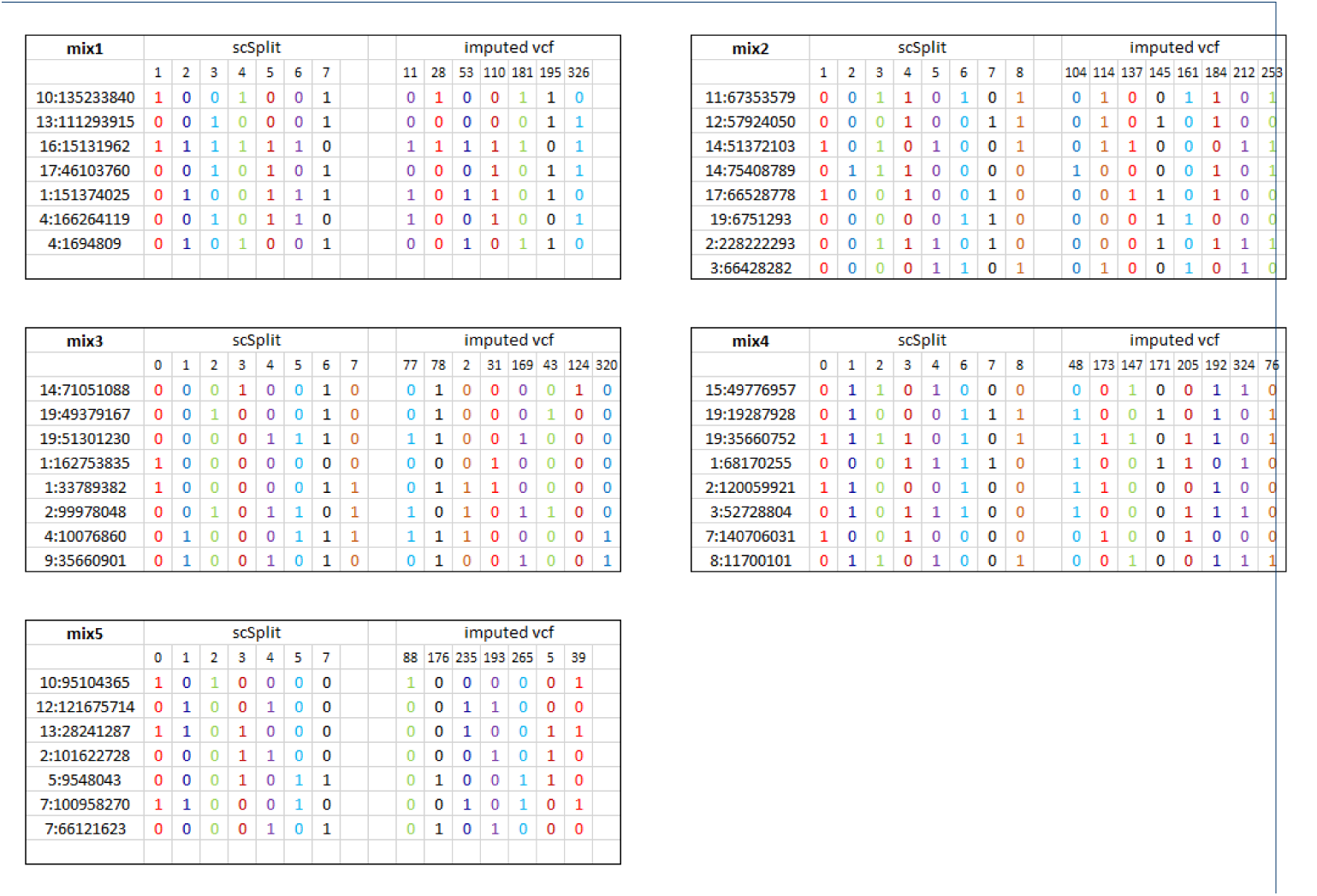
Illustration of presence absence matrices calculated on pooled scRNA-seq datasets. Within each rectangle we show the P/A genotype for each sample caclulate on scRNA-Seq data using scSpit, and the corresponding P/A genotype calculated from matched genotype files. The mapping of scRNA-Seq sample to genotype sample is indicated by colors.

